# Accounting for unobserved spatial variation in step selection analyses of animal movement via spatial random effects

**DOI:** 10.1101/2023.01.17.524368

**Authors:** Rafael Arce Guillen, Finn Lindgren, Stefanie Muff, Thomas W. Glass, Greg A. Breed, Ulrike E. Schlägel

## Abstract

1. Step selection analysis (SSA) is a common framework for understanding animal movement and resource selection using telemetry data. Such data are, however, inherently autocorrelated in space, a complication that could impact SSA-based inference if left unaddressed. Accounting for spatial correlation is standard statistical practice when analyzing spatial data, and its importance is increasingly recognized in ecological models (e.g., species distribution models). Nonetheless, no framework yet exists to account for such correlation when analyzing animal movement using SSA.
2. Here, we extend the popular method *Integrated Step Selection Analysis* (iSSA) by including a *Gaussian Field* (GF) in the linear predictor to account for spatial correlation. For this, we use the Bayesian framework `R-INLA` and the *Stochastic Partial Differential Equations* (SPDE) technique.
3. We show through a simulation study that our method provides unbiased fixed effects estimates, quantifies their uncertainty well and improves the predictions. In addition, we demonstrate the practical utility of our method by applying it to three wolverine (*Gulo gulo*) tracks.
4. Our method solves the problems of assuming spatially independent locations in the SSA framework. In addition, it offers new possibilities for making long-term predictions of habitat usage.

## 1 INTRODUCTION

Fine-scale animal tracking has become a popular tool in ecology and conservation research (Kays et al. 2015, Nathan et al. 2008). An important application of animal movement data is to make inference about the effects of environmental resources on movement decisions of animals (Thurfjell et al. 2014). For instance, Marshall et al. (2020) analyzed telemetry data of King Cobras in Northeast Thailand based on different landscape types. They showed evidence that these animals tend to move less on agricultural landscapes. Similarly, Prokopenko et al. (2017), studied the influence of roads on animal movement behaviour. Their analysis suggests that crossing roads was avoided by elk, even when traffic was low. Understanding these mechanisms is essential, as it allows humans to identify how animals react to disturbance elements or identify important habitat features, crucial for effective management and conservation. From a basic science perspective, these methods allow ecologists to understand crucial processes such as species distributions, home range formation, and assessing the intensity of species interactions (Fortin et al. 2005, Matthews et al. 2020, Thurfjell et al. 2014).

Among the most commonly used statistical tools for analysis of animals’ habitat selection using telemetry data are *resource selection functions* (RSF) (Boyce et al. 2002, Manly et al. 2007) and *step selection functions* (SSF) (Fortin et al. 2005, Forester et al. 2009). The main aim of these approaches is to understand which landscape features or different habitat types influence animal space use and to quantify the intensity of affinity or aversion to these explanatory variables. Methodologically, the original idea is to compare observed animal locations with a sample of locations that were available to the animals (Lele et al. 2013). For RSF, available locations are typically sampled uniformly over an area that is, in principle, deemed accessible to the studied animals, and the common practice is to use a logistic regression to make statistical inference on the parameters of the RSF. An inherent assumption of this approach is that there are no movement constraints for animals, i.e the animals could reach any location of the study area between consecutive observation times (Fieberg et al. 2021). However, this is not plausible if the time interval between consecutively observed locations is not large enough to assume their independence, which is almost always the case when RSFs are used to analyze movement data collected by modern biotelemerty devices (Nathan et al. 2022). To better account for this, SSFs instead make inference about consecutive steps that connect sequential locations rather than treating each location as independent, implemented through conditional logistic regression (Forester et al. 2009, Fieberg et al. 2021). Over the years, the use of SSFs has been refined, including different approaches to sample available steps (Forester et al. 2009), a correction for methodological approximations (iSSA, Avgar et al. 2016), and a reformulation of the conditional logistic regression approach as Poisson GLM to account for individual-level effects on habitat selection strength (Muff et al. 2020). Nowadays, many data sets are modelled with help of SSFs (also termed *step selection analysis*, SSA) rather than RSF approaches. A more detailed summary of the development of SSA can be found in Northrup et al. (2022).

Despite SSA being a well-established inference framework for telemetry data, there are some weaknesses. The SSA framework assumes steps to be spatially uncorrelated to each other since it does not fully account for the spatial nature of the data generating process. The true process is governed by complex behavioural decisions of animals in a heterogeneous landscape with potentially many influencing environmental factors. Since rarely all factors influencing movement known are measured, the unexplained spatial variation leads to spatial correlation. Ignoring this can cause an underestimation of the uncertainty of the parameters of interest (Fieberg et al. 2021), and underestimation of uncertainly can lead to incorrectly inferring statistical significance. In addition, without accounting for unexplained spatial variation, estimates of the means of the fixed effects can be biased in unknown directions (Thaden & Kneib 2018, Dupont et al. 2020). In SSA approaches, the quality of the model depends exclusively on the covariates provided by the user. Examples of missing covariates that could strongly influence movement include home ranges of species or unobserved individuals that interact with tracked individuals, which are difficult to identify *a priori* and can have a considerable effect on space-use decisions (Börger et al. 2008; Noonan et al. 2019). Another example of unexplained spatial variation is the concept of the landscape of fear in the context of predator-prey relationships. The risk of predation may cause prey to avoid certain regions and influence their landscape-feature selection. However, the *landscape of fear* is rarely included in analyses since it is nontrivial to quantify (Gallagher et al. 2017; Gaynor et al. 2019; Gehr et al. 2017). Neither SSA nor the RSA in their current form have the flexibility to explain and compensate for spatial correlation caused by omitting relevant, but unobserved, explanatory spatial variables.

In a variety of modelling contexts, a common approach to account for spatial, temporal or spatiotemporal correlation is to incorporate a continuous-space Gaussian field (GF) into the linear predictor. A spatial GF is a random effect which follows a multivariate Gaussian distribution with mean zero and spatial dependent covariance matrix, which is typically dominated by two hyperparameters, the spatial range and the standard deviation. Based on a flexible combination of these hyperparameters, the GF can take different shapes and thus capture various patterns of spatial dependence in the data (Lindgren et al. 2011). Including a GF in statistical models plays a fundamental role in present spatial statistics (Gelfand & Schliep 2016, Lindgren et al. 2011, Lindgren et al. 2022) and is becoming popular in species distribution modelling (Renner et al. 2015, Ward et al. 2015, Lezama-Ochoa et al. 2020, Engel et al. 2022). Although such species distribution models are somewhat related to SSA models for telemetry data, the latter need to account for the additional sequential nature of the data. Perhaps due to this added difficulty, GFs have not yet been implemented in SSA models.

Incorporating random effects into SSA models has long been restrictive due to the lack of existing software to fit the resulting mixed conditional logistic regression model. Fortunately, as Muff et al. (2020) point out, the conditional logistic regression model is a special case of a multinomial model and thus, it is likelihood-equivalent to a conditional Poisson model (conditioned on the number of observed events being fixed). In addition, Aarts et al. (2012) showed in the context of RSA that the maximum likelihood estimators of the slope parameters of a conditional *non-homogeneous Poisson process* (NHPP) are equivalent to the ones from the unconditional non-homogeneous Poisson process. Thus, a conditional NHPP can be fitted as an unconditional Poisson model with strata-specific intercepts (Aarts et al. 2012). Using the Poisson specification puts us in the *Generalized Linear Models* framework and therefore, random effects can be naturally incorporated (Muff et al. 2020). However, Muff et al. (2020) did not include the movement kernel in their implementation. Here we combine the contributions of Aarts et al. (2012) and Muff et al. (2019) to show analytically how the movement kernel can be also included in the Poisson implementation. In addition, we extend this to make use of the spatial nature of telemetry data and to account for missing spatial variation, by adding a spatial GF to the SSA framework. We take advantage of the fact that incorporating a GF is nothing else than including spatial random effects in the model. Although conceptually straightforward, the implementation of this is more complex, since we are fitting a hierarchical spatial model.

A popular inference method in the Bayesian framework is using the *Integrated nested Laplace approximation* (INLA). In general, when fitting hierarchical models, INLA offers a faster alternative to *Markov chain Monte Carlo* (MCMC) approaches. Unlike MCMC sampling methods, INLA is based on deterministic approximations of the marginal posterior distributions of fixed- and random effects as well of hyperparameters (Rue et al. 2009). Here, we use the R package inlabru, which is based on INLA (Bachl et al. 2019). This package is specialized in dealing with spatially structured data and is particularly convenient when fitting point processes. Simpson et al. (2016) illustrate the benefits of using a mesh-based approach known as *Stochastic Partial Differential Equations* (SPDE) when fitting an unconditional NHPP with random intensity. For this reason, we use the SPDE approach from Lindgren et al. (2011). An unconditional NHPP with random intensity is known as a *Log Gaussian Cox Process* (LGCP) (Diggle et al. 2013). We show analytically that we can use an LGCP to incorporate spatial random effects into SSA models.

We name our model *Gaussian Field integrated Step Selection Analysis* (GF-iSSA). With our method, we model unexplained spatial variation in the data, allowing users to avoid violating the spatial independence assumption of the model and to make more reliable inference about the fixed effects. We demonstrate the utility of our approach by applying it to simulated data and real data from three female wolverines (*Gulo gulo*).

## 2 MATERIALS AND METHODS

### 2.1 Model description

Telemetry data consist of time series of animal locations *s_t_*. At each time point, we specify the probability to observe an animal at a location, given where we observed it at the previous two times. In the classical step-selection model (Forester et al. 2009), this is done based on two aspects:

i. The general movement tendency of an animal in the absence of habitat selection, modelled by a movement kernel *ϕ*, also termed selection-free movement kernel (Fieberg et al. 2021).
ii. Selection behaviour of the animal with respect to environmental variables, modelled by a RSF *ω*.

The spatial density of observing an animal at location *s_t_* at time point *t*, given the last two observed steps *s_t_*_−1_, *s_t_*_−2_ and a landscape ***X***, is then modelled as (Forester et al. 2009):

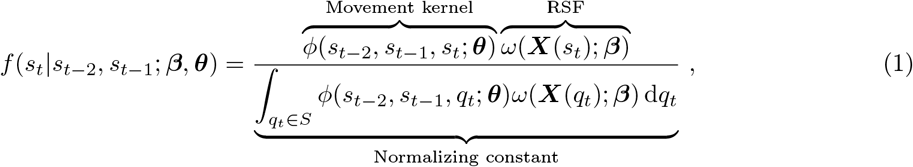

where *S* represents the spatial domain over which the animal may possibly move.

The selection function *ω* is modelled analogously to a RSF as:

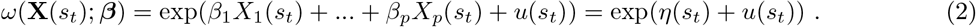

Here *X_i_*, *i* = 1, *…, n*, are spatial covariates, and we are interested in making inference about their effects on movement decisions. The term *u* represents a GF, which accounts for the spatial variation not explained by the fixed effects *η*(*s_t_*). Thus, *u* follows a multivariate Gaussian distribution with covariance matrix *C*:

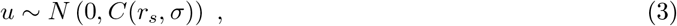

where the hyperparameters *r_s_* and *σ* represent the spatial range and the standard deviation, respectively.

The movement kernel *ϕ* is commonly modelled via a product of probability density functions for step lengths (SL) and turning angles (TA), and it depends on the parameter vector ***θ***. This requires a transformation from polar to euclidean coordinates through the change-of-variable technique, which results in an extra factor corresponding to the reciprocal of the step length (Schlägel & Lewis 2016). Similar to Avgar et al. (2016), we assume a gamma and a von Mises distribution for step lengths and turning angles, respectively. However, any distribution from the exponential family would work in our approach.

Note that the movement kernel *ϕ* depends on *s_t_*_−1_ for calculating the step lengths and additionally on *s_t_*_−2_ for the turning angles. Due to the special form of densities belonging to the exponential family, we can express the movement kernel as (Avgar et al. 2016, Munden et al. 2021):

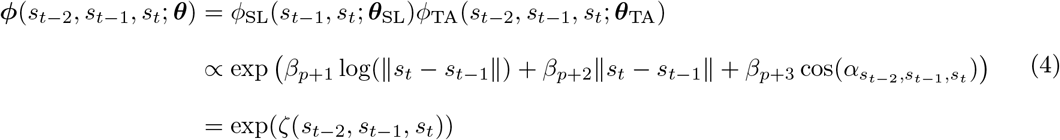

Here *β_p_*_+1_ and *β_p_*_+2_ are linked to the shape and rate parameter *a* and *b* of the gamma distribution, respectively. In addition, *β_p_*_+3_ represents the concentration parameter *k* of the zero-mean von Mises distribution. A detailed derivation of Eqn (4) can be found in the Appendix section **??**.

Combining the movement kernel (Eqn (2)) and selection function (Eqn (4)), we can express the likelihood (Eqn 1) as follows:

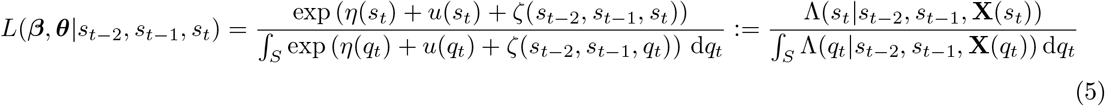

This is the likelihood function of a *conditional* non-homogeneous Poisson process (NHPP) (Aarts et al. 2012). Classic approaches to SSA first approximate the integral from Eqn (1) by sampling integration points from an empirically parameterized movement kernel *ϕ*^*^ based on the observed step lengths and turning angles (Forester et al. 2009, Fieberg et al. 2021). The resulting equation has consequently the form of a conditional logistic likelihood with discrete space. In contrast to this, we interpret telemetry data at each time point directly as an observation from a conditional NHPP. From a practical perspective, however, it is more convenient to implement an *unconditional* NHPP. This can be achieved by using at each time point an unconditional NHPP with an additional intercept. It has been shown that for such a model, the maximum likelihood estimator of the slope parameters are equivalent to the ones from the conditional NHPP (Aarts et al. 2012, Muff et al. 2020). This equivalence can also be shown in a Bayesian framework for the posterior distribution of our parameters (Appendix section **??**). Therefore, the joint log-likelihood of our model for a total of *T −* 2 time points (The first two locations are used to calculate the first TA and SL of the first step) results in the following:

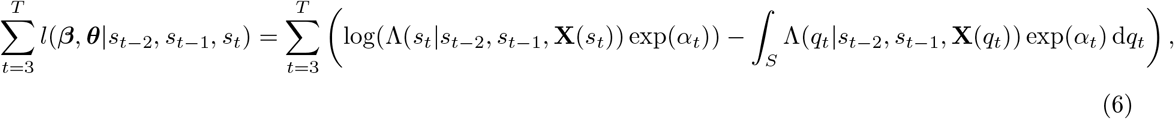

where *α_t_*, *t* = 3*,…, T* represent the time dependent intercepts which allow us to use the unconditional NHPP at each time point. Note that the intensity function Λ(*s_t_|s_t_*_−1_, *s_t_*_−2_, **X**(*s_t_*)) is stochastic due to the presence of spatial random effects *u*(*s_t_*), making it a hierarchical model. This flexible model is known as a *Log Gaussian Cox Process* (LGCP). Given a realisation of Λ(*s_t_|s_t_*_−1_, *s_t_*_−2_, **X**(*s_t_*)), at each time point the model is a NHPP (Diggle et al. 2013).

The interpretation of the parameters ***β*** of the selection function is the same as the one presented by Fieberg et al. (2021). For example, given two locations *s_u_* and *s_v_* that are equally available (based on the movement kernel), have the same spatial resources but only vary by one unit of *X*_1_, then the relative use of these two locations is equal to exp(*β*_1_) (Fieberg et al. 2021).

### 2.2 Model fitting

We performed Bayesian inference with the inlabru package (Bachl et al. 2019). To fit the model specified in Eqn (6) to data, the integral from Eqn (6) was computed numerically over discretized space. Using the SPDE method, the model needs an approximation of the GF. This is achieved via linear basis functions applied at integration points defined by all the nodes of the mesh (Simpson et al. 2016). The mesh is a triangulation of the domain (Lindgren et al. 2011). The purpose of this is to capture all the spatial effects not included in the fixed effects. We therefore approximated Eqn (6) by:

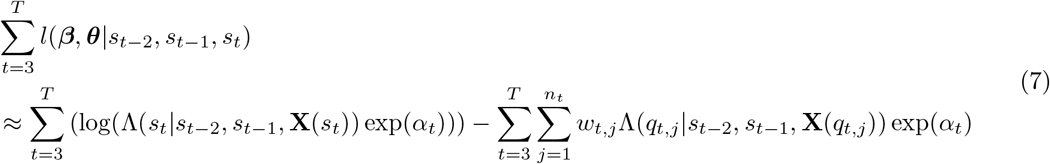

for suitable integration points *q_t,j_* and weights *w_t,j_*. For this, we defined a uniform mesh over the study area (Fig. 1) and used the nodes of the mesh both as integration points and to approximate the GF by a Gaussian Markov Random Field (Simpson et al. 2016). Thus, the model can be reformulated to a single Poisson model with time-specific intercepts. Since we are not interested in making inference about them and for numerical reasons, we apply the solution of Muff et al. (2020) and specify them as random intercepts with a large fixed variance in order to avoid shrinkage.

The use of a deterministic integration scheme instead of a random *Monte Carlo* integration approach has two motivations. First, the integration scheme needs to capture the full range of freedom of the random field components in order to give a valid approximation of the likelihood. And second, for spatial dimensions lower than 4, the numerical bias in a basic deterministic integration scheme decreases faster than the standard deviation of a Monte Carlo integration scheme, for an increasing number of integration points. By placing an integration point at each mesh node, both aspects are taken into account.

**Figure 1:**
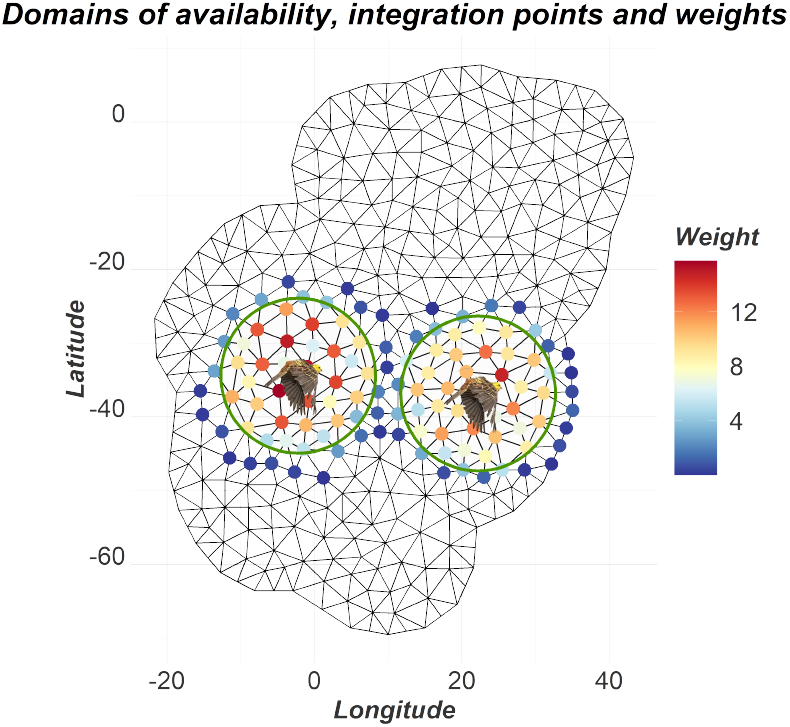
Domains of availability for two time points. The birds represent the observed animal locations. The nodes of the mesh are used as integration points with their corresponding weights based on linear basis function integrals.

To reduce computational burden, we restricted the domain of availability *S* and hence integration points at each time point to a disk around the observed location *s_t_* with a radius at least equal to the maximum observed step length over the entire data set (Fig. 1). This means that we essentially truncated the kernel *ϕ* at a distance for which the probability of observing a step became negligible. The integration points serve a similar purpose as the available steps in the classical SSA framework, i.e. computing numerically an approximation to the integral from Eqn (1). The weights were calculated based on the integrals of the piecewise linear basis functions used to represent the spatially discretised GF model. This generated non-zero weights for all triangle vertices of triangles that intersect the corresponding availability disk (Fig. 1), computed via dense deterministic sampling within each triangle (Jullum 2020). This step was automatically performed with help of the inlabru package. A code example of our approach can be found in the supplementary material.

For all parameters of the selection function and movement kernel, we used the default `R-INLA` uninformed priors, i.e. a Gaussian distribution with mean equal to zero and a large variance. We opted for this since for real data applications, we prefer to relax the assumption of the gamma and von Mises distribution by letting the parameters be estimated without constraints. Furthermore, we used the penalized complexity priors for our hyperparameters as explained in Gómez-Rubio (2020). Consequently, the penalized complexity priors are defined as probability beliefs:

1. 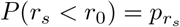
2. 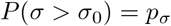

(Fuglstad et al. 2019). Thus, the users can assign their prior beliefs through *r*_0_, 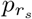, *σ*_0_ and *p_σ_* respectively. Notice that *r_s_* represents the practical range, which is the distance at which the spatial correlation is around 0.139 (Krainski et al. 2018). When setting the probabilities 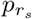 and *p_σ_* equal to 0.05, Fuglstad et al. (2019) suggest to assign a value *σ*_0_ which is 2.5 to 40 times the true value of the standard deviation. In addition *r*_0_ can be set to a value which should be between 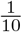 and 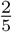 of the true range. This specification leads to stable inference results of the marginal posterior distributions (Fuglstad et al. 2019).

Note that since this method uses Bayesian and not a frequentist inference, classical model comparison with help of the Akaike information criterion (AIC) is not possible. However, R-INLA provides a Bayesian alternative to the AIC called the deviance information criterion (DIC). Similar to the AIC, a model with a lower DIC should be preferred (Gómez-Rubio 2020).

### 2.3 Simulation study

We simulated animal tracks based on the step-selection model defined by the conceptional likelihood from Forester et al. (2009) (Eqn (1)). For this, we discretized space, using a fine grid over the whole study area represented by a raster object of resolution 1000 *×* 1000 and assigning at each time point a likelihood value for each grid cell.

For the movement kernel, we specified a gamma distribution for the step lengths with shape and rate parameters equal to 5 and 2 respectively. In addition, we specified for the turning angles a von Mises distribution centered at 0 with concentration parameter set to 1. Thus, these animals had a slight tendency to move in a straight direction.

For the selection function, we used three fixed effects: A continuous variable x1, a discrete covariate x2 and the distance to home range center cen. Their corresponding selection coefficients were set to 1.5, 1 and *−*0.04. The variable *cen* implements a centralizing tendency that accounts for home ranging behaviour of animals (Fig. 2). In real applications x1 could for example represent elevation and x2 could describe a variable representing different landscape types. In addition, we included as a raster layer a sample of a GF with a Matérn covariance function with help of the geoR package (Ribeiro Jr et al. 2020) and incorporated it in the linear predictor. The GF represents the spatial variation not explained by the fixed effects. We simulated in total nine scenarios using different combinations of spatial ranges *r*_true_ (30, 40, 50) and standard deviations *σ*_true_ (1, 2, 3) hyperparameters for the GF. The geoR package was also used to generate the environmental covariates. For each scenario, we simulated 25 animal tracks leading to 225 animal tracks. Each track consists of 1000 observed locations. We made sure that there is no high correlation (*|ρ|*< 0.10) between the GF and the fixed effects to avoid confounding results.

**Figure 2:**
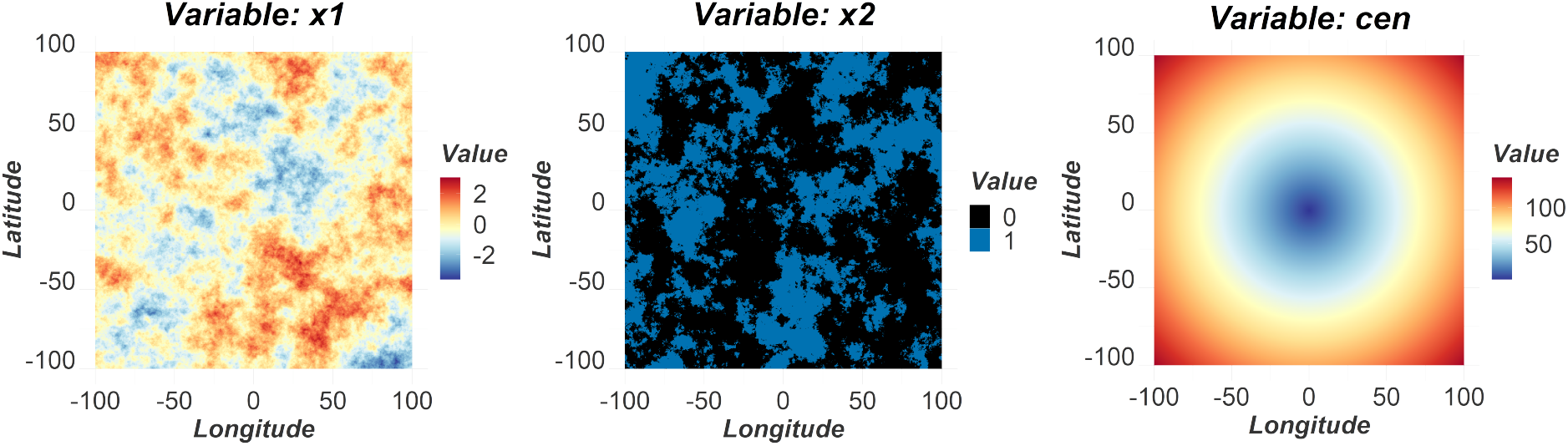
Environmental covariates of the simulation. Continuous covariate *x*1 (left), discrete covariate *x*2 (middle) and distance to home range center *cen* (right).

We fit separate models to each individual track. In order to fit the models, the radius of the domain of availability at each time point was set to 1.5 times the maximum observed step length. Since the theoretical mean of the step length distribution is equal to 2 spatial units, we preferred to used a mesh resolution lower than this. Thus, we defined our meshes to have a maximum edge length of 1.5 spatial units. For the hyperparameters of the GF, we used the penalized complexity priors as follows:

1. 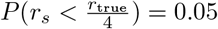
2. 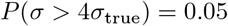

The choice of these priors was based on Fuglstad et al. (2019) and thus, other prior choices would work properly as well.

To compare our new approach to the most commonly used one, we also performed iSSA (Avgar et al. 2016) on all simulated tracks, using two approaches. First, we fitted the model with the sampled GF used in the simulations as a known covariate, i.e. we treated the GF as a fixed effect (Full-iSSA). This scenario is, however, utopian for real data applications. Second, in order to observe how the iSSA reacts to missing spatial covariates, we applied the iSSA method excluding the GF and compared the results with our GF-iSSA approach. Both the Full-iSSA and the iSSA were fitted using a conditional logistic regression. For this, we sampled 500 available locations per used location from the initial movement kernel parametrized based on the observed step lengths and turning angles using the R package amt (Signer et al. 2019). We compared the mean estimates of the fixed effects of all three models. In addition, since the general purpose of making inference is quantifying uncertainty, we calculated the coverage properties of the GF-iSSA in comparison to the iSSA approach. Thus, for all simulations, we calculated for each fixed effect the rate for which the true value was covered by the corresponding 95% credible intervals and confidence intervals, respectively. Finally, we investigated graphically how the contribution of the GF can increase the predictive quality of the model.

### 2.4 Case study

As exemplary case study, we applied our method to GPS collar data from three female wolverines in Arctic Alaska (Glass et al. 2021). The data were collected between March 2017 and February 2019 in the vicinity of Toolik Field Station (68.63° N, 149.60° W). Wolverines were equipped with GPS collars, programmed to obtain location coordinates every 40 minutes. For more details, see Glass et al. (2021).

We here used as spatial covariates three of the environmental variables from the habitat selection analysis of Glass et al. (2021): (i) terrain ruggedness index, both as linear and quadratic term to account for a possible non-linear relationship (ii) distance to streams and rivers, and (iii) distance to lake edges. Glass et al. (2021) also included snow depth and density as spatiotemporal explanatory covariates. We omitted these variables in our analysis and instead tested the GF’s ability to adjust for them and any other missing information. Similar to Glass et al. (2021), all covariates were standardized. We have applied the iSSA and the GF-iSSA to three female wolverine tracks “F6", “F11” and “F12”. The Full-iSSA could not be applied here since the true data generating process is unknown. Given that we observed some large outliers in the observed step lengths, we defined the ratio for the domain of availability as the 0.95-quantile of the observed step lengths. All other model fitting specifications were analogous to the simulation study.

## 3 RESULTS

### 3.1 Simulation study

Overall, the GF-iSSA reliably estimated the posterior means of the fixed effects. The estimates of the selection coefficients were on average unbiased (Fig. 3). The selection coefficients posterior mean estimates of our method deviated absolutely on average by 0.11 from the simulated values. In addition, our approach seemed not to be overconfident given that the box-plots cover in general the true values. This was also true for the movement kernel parameters, for which our model performed well despite the presence of missing spatial variation (Fig. 3). Here the average absolute deviation from the true movement kernel parameters was 0.08. In seven out of the 225 animal tracks our method failed or crashed numerically, presumably due to unsuitable prior specification of the GF. Therefore, the results include 218 animal tracks. We calculated the running times for the GF-iSSA approach. Our models had a median running time of around 29 minutes per track. The 25%-quantile and the 75%-quantile were equal to around 24 and 37 minutes. However, we had an outlier with 15.5 hours. We used a Debian GNU/Linux 11 cluster specifying two cores for each model.

**Figure 3:**
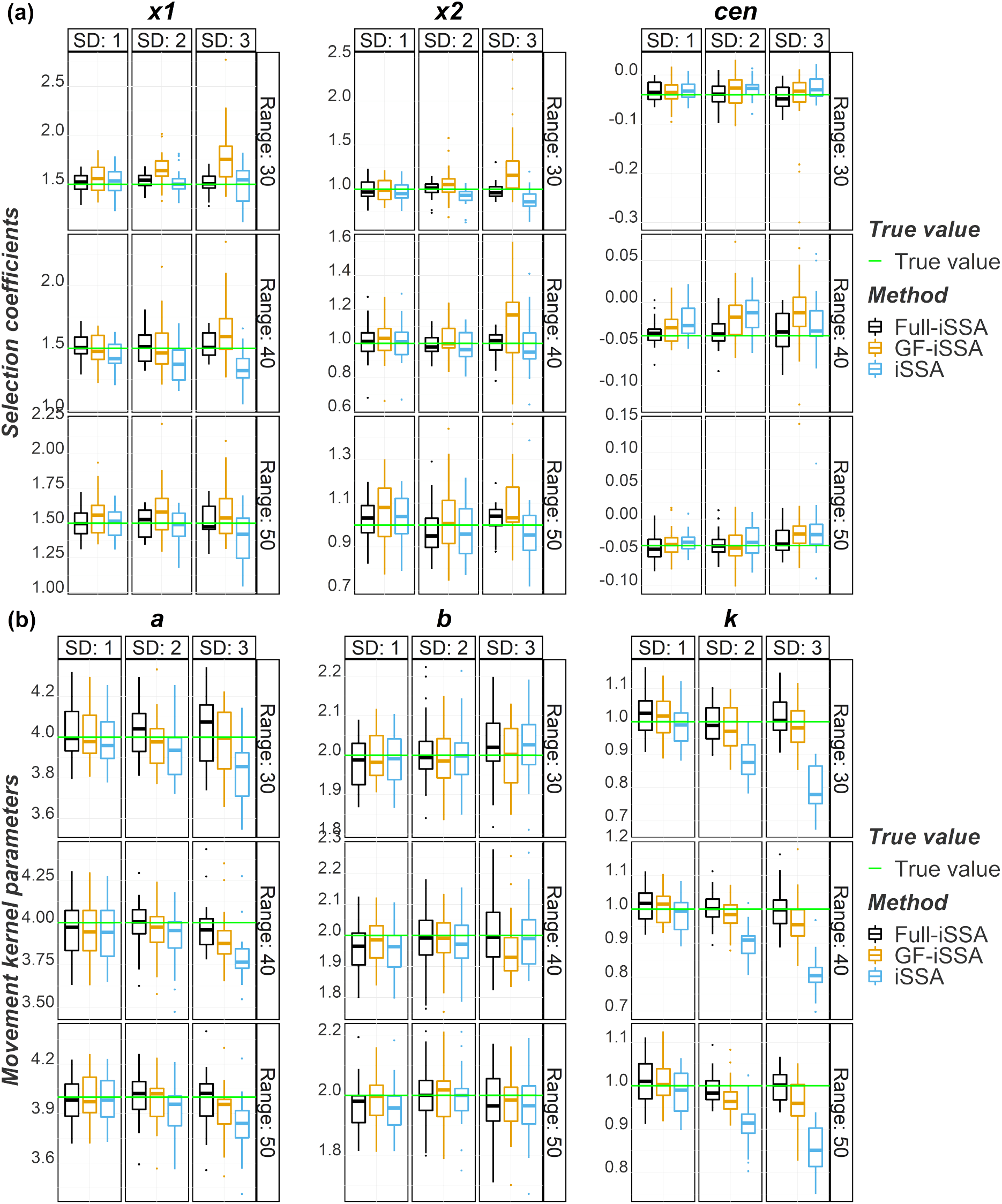
Mean estimates of fixed effects in the simulation study: GF-iSSA vs iSSA. Comparison of (a) selection coefficients for the three covariates x1, x2, and cen, and (b) movement kernel parameters, for various combinations of the hyperparameters (standard deviation and spatial range) of the simulated Gaussian field. The black box-plots represent the Full-iSSA method. The yellow box-plots represent the GF-iSSA and the blue box-plots represent the iSSA. The latter does not account for the missing spatial variation. The green horizontal lines represent the true values of the parameters.

When comparing the GF-iSSA to the iSSA approach, we found that in general both methods returned similar mean estimates in case of the environmental parameters. However, the GF-iSSA showed in general wider box-plots than the iSSA suggesting that the GF-iSSA is more conservative in presence of spatial autocorrelation (Fig. 3). When the standard deviation of the GF was low (SD = 1), both methods estimated the coefficients stably. However, when the standard deviation hyperparameter increased, both approaches seemed to show slightly more variability across the estimated selection coefficients compared to the Full-iSSA. A difference between our method and the iSSA was that the GF-iSSA tended to sometimes overestimate the selection strength while iSSA occasionally underestimated it. In general both methods estimated the mean estimates of the selection coefficients decently under missing spatial variation.

The mean estimates of the movement kernel parameters were, however, noticeably less biased in case of the GF-iSSA. Here we could also clearly observe that the estimates of the iSSA were systematically underestimated with an increasing standard deviation of the GF while the ones from the GF-iSSA remained more centered to the corresponding true values (Fig. 3). This was true for the shape parameter *a* and the concentration parameter *k*. Despite being unbiased, the box-plots did not show a large variation suggesting that the iSSA is overconfident at estimating these parameters. As expected, the utopian Full-iSSA remained stable and unbiased for all the fixed effects.

On average, for all the simulations of the nine scenarios, both the iSSA and GF-iSSA returned decent mean estimates of the fixed effects, which did not deviate significantly from the true values (Table 1). However, the posterior mean estimates of the GF-iSSA showed slightly larger variation compared iSSA estimates based on the 5%- and 95% quantile of the mean estimates. In addition, the coverage results of the GF-iSSA were better than the ones from the iSSA. For all the fixed effects with the exception of the coefficient for *x*2, our method had a higher coverage than the iSSA approach. In addition, our method had a mean coverage of around 91% while the iSSA had a mean coverage of 82% (Table 1), meaning that in presence of missing spatial variation the uncertainty of the iSSA parameters was underestimated compared to the uncertainties estimated from the GF-iSSA, which was closer to the expected 95% coverage.

**Table 1:** Mean estimates of fixed effects. The results of each parameter are based on 218 tracks, summarized from all nine simulation scenarios. The quantiles are those of the mean estimates. The coverage represents the percentage for which the corresponding true values were covered by the 95% credible intervals (GF-iSSA) and Wald confidence intervals (iSSA) respectively.

| Parameter | Method | Mean | True value | 0.05-quantile | 0.95-quantile | Coverage(%) |
| --- | --- | --- | --- | --- | --- | --- |
| x1 | Full-iSSA | 1.51 | 1.50 | 1.35 | 1.68 | 0.95 |
| x1 | GF-iSSA | 1.60 | 1.50 | 1.31 | 1.95 | 0.82 |
| x1 | iSSA | 1.45 | 1.50 | 1.18 | 1.70 | 0.76 |
| x2 | Full-iSSA | 1.00 | 1.00 | 0.85 | 1.15 | 0.96 |
| x2 | GF-iSSA | 1.07 | 1.00 | 0.82 | 1.44 | 0.82 |
| x2 | iSSA | 0.97 | 1.00 | 0.77 | 1.17 | 0.85 |
| cen | Full-iSSA | -0.04 | -0.04 | -0.07 | -0.00 | 0.96 |
| cen | GF-iSSA | -0.03 | -0.04 | -0.08 | 0.02 | 0.97 |
| cen | iSSA | -0.03 | -0.04 | -0.06 | 0.01 | 0.89 |
| a | Full-iSSA | 4.00 | 4.00 | 3.76 | 4.25 | 0.97 |
| a | GF-iSSA | 3.96 | 4.00 | 3.72 | 4.23 | 0.95 |
| a | iSSA | 3.90 | 4.00 | 3.61 | 4.19 | 0.91 |
| b | Full-iSSA | 1.99 | 2.00 | 1.84 | 2.15 | 0.93 |
| b | GF-iSSA | 1.98 | 2.00 | 1.84 | 2.12 | 0.95 |
| b | iSSA | 1.99 | 2.00 | 1.84 | 2.14 | 0.94 |
| k | Full-iSSA | 1.01 | 1.00 | 0.93 | 1.10 | 0.94 |
| k | GF-iSSA | 0.99 | 1.00 | 0.90 | 1.07 | 0.93 |
| k | iSSA | 0.90 | 1.00 | 0.75 | 1.04 | 0.55 |

The posterior mean estimates of the hyperparameters (parameters of the Gaussian field) showed a large variation and were often not centred at the true value (Fig. S1). However, we simulated the Gaussian field for the whole study area and not for the effective study area, which was the one after joining the domains of availability with help of the aforementioned disks. Thus, the true parameters were not identifiable.

The inlabru package can easily provide a map of the GF. The GF-iSSA approach estimated the GF well. It was able to detect the high and low impact areas and therefore effectively account for missing spatial covariates. We show illustratively the true GF, the estimated posterior mean distribution of the GF and a sample of it for one of our animal tracks (Fig. 4). In addition, the GF can have a large contribution to the linear predictor where missing covariates have a large effect on animal movement. When displaying the contribution to the linear predictor of the spatial covariates without the GF, the estimated selection function did not match the observed animal locations. However, when additionally including the GF this mismatch became much lower (Fig. 4).

**Figure 4:**
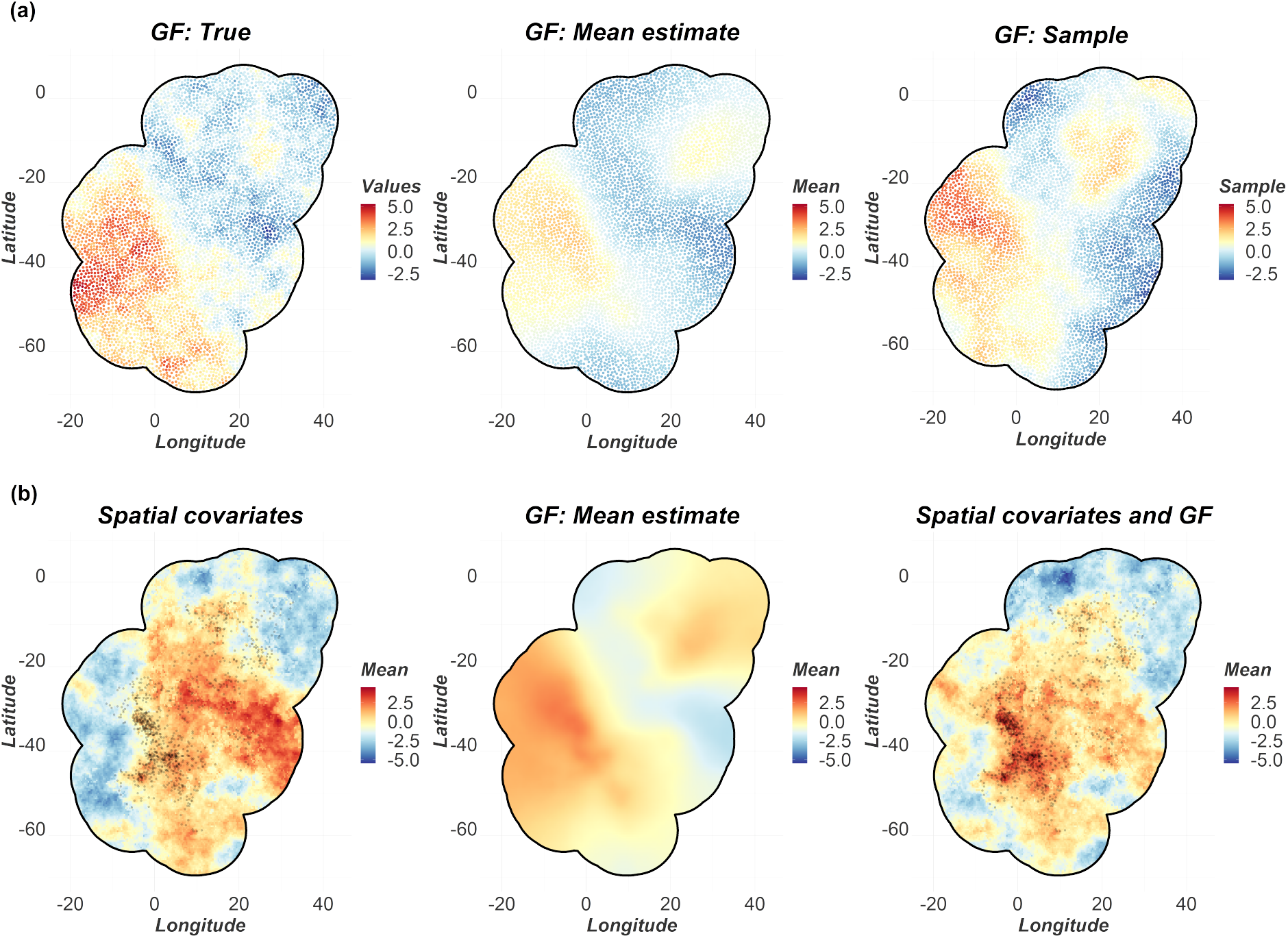
Visualization of the GF for one track and its predictive contribution. (a) Comparison between the true GF (left), the mean estimates of the GF (middle) and a sample of it (right). (b) Contribution to the linear predictor of the spatial covariates (left), the estimated GF (middle) and the combined effect (right). The black dots represent the observed animal locations.

### 3.2 Case study

The results of the GF-iSSA applied to the three wolverine tracks were satisfactory and demonstrate the utility of the GF-iSSA. Despite our analyses not including snow layers as covariates, which were known from Glass et al. (2021) to influence movement of wolverines, our model was able to estimate the selection coefficients plausibly. Our mean estimates were more in line with the original results from Glass et al. (2021) than the iSSA without the snow layers presented here (Table 2). Based on the reported credible intervals, for all three wolverine tracks, there was strong statistical evidence for a positive effect for terrain ruggedness (Table 2). In addition, our results report a negative selection for the squared terrain ruggedness indicating that at some point a high terrain ruggedness is avoided, although, based on the credible intervals, there was no strong statistical evidence for this effect in case of “F11” and “F12" (Table 2). Combining the effect of the linear and squared term, “F6” is more likely to avoid very low and high terrain ruggedness. In contrast, the estimated combined effect of the iSSA would suggest a monotonically increasing selection for high terrain ruggedness, which is implausible and not in line with Glass et al. (2021). In addition, our model returned negative posterior mean estimates for the distances to rivers. In all three cases, zero was not covered by the corresponding 95% credible intervals indicating a strong statistical evidence for an affinity to riparian habitat. With the GF-iSSA we could only detect strong statistical support for the distance to lakes in the case of “F6”. This was not true for the iSSA approach, which returned a positive estimate for lakes for “F6", again deviating from the results of Glass et al. (2021). However, there was little or no statistical evidence for this effect.

**Table 2:** Estimates of fixed effects for movement tracks of three female wolverines. Selection was estimated with respect to a terrain ruggedness index (terrain), distance to rivers and streams (rivers), and distance to lakes (lakes).

|  |  | iSSA |  |  | GF-iSSA |  |  |
| --- | --- | --- | --- | --- | --- | --- | --- |
| ID | Parameter | Estimate | CI_0.025 | CI_0.975 | Estimate | CI_0.025 | CI_0.975 |
| F6 | terrain | 0.49 | 0.38 | 0.60 | 0.20 | 0.18 | 0.22 |
| F6 | terrain<sup>2</sup> | -0.00 | -0.03 | 0.02 | -0.03 | -0.03 | -0.02 |
| F6 | rivers | -0.24 | -0.31 | -0.18 | -0.28 | -0.29 | -0.26 |
| F6 | lakes | 0.05 | -0.12 | 0.22 | -0.56 | -0.60 | -0.53 |
| F6 | a | 0.00 | 0.00 | 0.00 | -0.00 | 0.00 | 0.00 |
| F6 | b | 0.31 | -0.00 | 0.03 | 1.00 | -1.02 | -0.99 |
| F6 | k | 0.02 | -0.19 | -0.07 | -0.04 | -0.05 | -0.04 |
| F11 | terrain | 1.02 | 0.82 | 1.22 | 1.79 | 1.64 | 1.93 |
| F11 | terrain<sup>2</sup> | -0.47 | -0.76 | -0.18 | -0.06 | -0.16 | 0.04 |
| F11 | rivers | -0.95 | -1.15 | -0.75 | -0.33 | -0.48 | -0.18 |
| F11 | lakes | -0.02 | -0.30 | 0.25 | 0.07 | -0.19 | 0.32 |
| F11 | a | 0.00 | 0.00 | 0.00 | -0.00 | 0.00 | 0.00 |
| F11 | b | 0.29 | 0.01 | 0.05 | 0.31 | -1.74 | -1.65 |
| F11 | k | 0.02 | -0.33 | -0.11 | -0.48 | -0.58 | -0.38 |
| F12 | terrain | 1.27 | 0.98 | 1.56 | 2.48 | 2.10 | 2.85 |
| F12 | terrain<sup>2</sup> | 0.00 | -0.31 | 0.31 | -0.16 | -0.47 | 0.15 |
| F12 | rivers | -1.06 | -1.40 | -0.72 | -1.01 | -1.39 | -0.62 |
| F12 | lakes | -0.08 | -0.32 | 0.15 | -0.12 | -0.42 | 0.18 |
| F12 | a | 0.00 | -0.00 | 0.00 | -0.00 | 0.00 | 0.00 |
| F12 | b | 0.38 | 0.01 | 0.11 | -1.75 | -3.97 | -3.52 |
| F12 | k | 0.41 | 0.04 | 0.38 | 0.08 | -0.12 | 0.29 |

The effect of the unexplained spatial variation was, however, low for the wolverine data. For our three tracks, the GF-iSSA estimated on average a spatial range of 1249 meters and a standard deviation of 0.28. Thus, given the low standard deviation, the missing spatial variation is low compared to the contribution of the fixed effects in this case indicating that the spatial autocorrelation was low for these animals and that most of the spatial variation was explained by the fixed effects.

Note that the movement kernel estimates from our approach differed from the iSSA model. Interpreting the parameters from the GF-iSSA as the theoretical parameters of the gamma and von Mises distributions, we would obtain negative scale *a*, shape *b* and concentration *k* parameters (Table 2), due to the model not accounting for the fact that these parameters are theoretically bounded. However, when displaying the estimated combined effect of SL and log(SL), these parameters produced a sensible movement kernel similar to a gamma distribution and not deviating significantly from the observed step lengths. In addition, the negative concentration parameter indicated that the mean of the von Mises distribution was not necessarily 0, as expected for data relatively large time intervals between consecutive locations (here every 40 minutes).

## 4 DISCUSSION

Here we introduced an approach for incorporating and estimating spatial random effects in step selection analyses. This method, which we termed GF-iSSA, helps to resolve a perennial problem in animal movement analyses by accounting for spatial autocorrelation arising from unobserved spatial effects in habitat selection analyses via a continuous-space Gaussian field. Rather than being an alternative model, the GF-iSSA can be interpreted as an extension of the iSSA, which is able to account for missing spatial covariates. Incorporating random effects leads to better estimates of fixed effects parameters, which in current SSA frameworks may be contaminated by unaccounted spatial covariates. In addition, the estimate of the Gaussian field itself can be biologically interpreted and help to identify hitherto unknown causes of spatial autocorrelation in observed movement (e.g. landscape features). We have shown that our approach works on both simulated and empirical data. Our model estimated fixed effects means and uncertainties reliably. Including spatial random effects in analyses of spatially structured data has become a common practice in the last decade; analyses of animal movement data should not be an exception.

Based on a simulation study, we have shown that the GF-iSSA reliably estimates fixed-effects for various simulation settings. Overall, our method produced nearly unbiased estimates for both the selection coefficients and the movement kernel parameters. In particular, GF-iSSA estimates of the movement-kernel parameters were consistently close to the underlying simulated parameter values, in contrast to estimates from iSSA, which showed a tendency to underestimate parameters. Regarding the habitat selection parameters, the difference in mean estimates from GF-iSSA and iSSA was less pronounced, showing some robustness of the iSSA under missing spatial variation. In general, for both the GF-iSSA and the iSSA, variability in the mean estimates increased as the standard deviation of the GF increased. However, not accounting for spatial correlation (i.e. using iSSA) resulted in an underestimated uncertainty of these parameters. Thus, based on the coverage results, the GF-iSSA quantifies uncertainty more reliably and is therefore more likely to prevent underestimating the uncertainty and resultant increased risk of spurious statistical support. In addition, our method estimated the GF accurately and is therefore able to represent the underlying spatial process in its entirety. In case of the three wolverine tracks, the GF-iSSA recovered the parameters of the static habitat features from the original analysis of Glass et al. (2021), even though we omitted the spatio-temporal snow layers. This indicates that the GF-iSSA could explain most of the spatial variation of the missing snow covariates, allowing for reliable inference of the fixed effects that were included.

The case study revealed that using deterministic integration points via a mesh and therefore estimating the parameters of the movement kernel directly could lead to estimates that are outside their theoretical domains (e.g. negative values for the shape or rate parameter of a Gamma distribution). Nonetheless, this would also be the case if the user fit the iSSA with deterministic integration points. Thus, this problem does not arise from the added GF, but rather from the integration strategy. In addition, although less likely, the same problem may also occur even when sampling integration points from an initial movement kernel. Accounting for this model behaviour with restrictive priors led to numerical issues that prevented model fitting. Thus, we opted to allow the movement kernel parameters to be estimated without constraint. Enforcing parameter space constraints, in addition to introducing numerical issues, may also result in incorrect inference. Despite all presented advantages of the GF-iSSA, it comes with a higher computational cost than some conventional alternatives, depending upon the desired resolution of the mesh. A finer mesh requires more integration points and commesurately longer computation times for model fitting. Fitting LGCPs is generally computationally expensive, but the aforementioned advantages, in our opinion, justify the relatively modest numerical expenditures here (average fitting time is about 30 minutes). In addition, for real data applications, the numerical stability of our method is sensitive to the choice of the priors for the hyperparameters of the GF.

For our simulation study, we assumed that the GF is independent of the fixed effects. This is a strong assumption, since in real situations the spatial effects represented by the GF are likely to affect both the dependent variable and the observed spatial covariates simultaneously. If this assumption is not met, this could lead to biased inference results given the potential high correlation between the GF and some spatial covariates (Thaden & Kneib 2018). This problem, commonly known as spatial confounding, however holds for both the iSSA and the GF-iSSA. A future avenue for development that might resolve this issue could be to use flexible spatial confounding approaches like *Spatial+* from Dupont et al. (2020) or the *Geoadditive Structural Equation Model* approach from Thaden & Kneib (2018). These approaches require first accounting for spatial autocorrelation in the linear predictor. Thus, the GF-iSSA opens new horizons for addressing this non-independence. Nonetheless, it is not yet clear how to adapt and formally implement these approaches in a Bayesian framework.

Finally, we emphasize that our model, as well as other SSA framework models, do not make inference about the true movement process of the animals since the data are discrete in time. Rather, it describes the observational process of animals at constant, discrete time intervals. However, methods using non-constant time intervals like the *time-varying iSSA* (tiSSA) from Munden et al. (2021) could be extended to include a GF as a spatial random effect in the linear predictor. For this, the domains of availability could be defined as disks where the radius equals the time difference multiplied by the maximum observed speed. In addition, our method can also be applied for making inference about tracks of multiple animals simultaneously using random effects as proposed by Muff et al. (2020).

For the future, it would be interesting to additionally include temporal autocorrelation in the covariance matrix of the GF. However, there is no purely temporal correlation in telemetry data, since the data’s time dimension is controlled by the tracking technology and not by the animal itself. Thus, the model presented here could only be extended using a non-separable spatiotemporal correlation matrix instead of the purely spatial Matérn correlation matrix. However, this is a numerical challenge. Another modification of the model could be to use circular time instead of linear time. Circular time is convenient when making inference about patterns in a repeating time window. In our case, the time window could be a 24-hours diel cycle, such as that used by Shirota & Gelfand (2017). Although their model was used for crime data, a similar approach could be used for telemetry data. For instance, Benoit-Bird et al. (2009) analysed the effect of nocturnal light on the diel migration of micronekton in the water column. In our case, the model of Shirota & Gelfand (2017) could account for the missing sunlight covariate, supposing that the sunlight had an impact on movement decisions, but the users do not dispose of this variable.

In summary, our study demonstrates how to use the GF-iSSA to account for unobserved spatial effects in habitat selection analyses. This approach has three advantages. The first and largest advantage is an appropriately estimated uncertainty, which is key to correct biological inference. Second, based on our simulation study, the mean estimates of the movement kernel are less biased including a GF in the model. Nonetheless, the selection coefficients estimates do not deviate meaningfully from those fitted using a normal iSSA. However, we recommend the users to focus not only on the mean estimates but also on their uncertainty, which in our simulation study was better estimated by GF-iSSA than iSSA. Third, although not the main focus of this paper, our method has a high predictive quality compared to methods not accounting for spatial autocorrelation. Via the Gaussian field, the GF-iSSA formally estimates unobserved spatial random effects, and can use these effects in addition to observed fixed effects to make predictions. Users additionally interested in predictions (rather than inference regarding the effects of particular habitat types on movement and space use) would thus benefit from the GF-iSSA. Although not representing the whole linear prediction, since the movement kernel is not included, the selection function of the GF-iSSA and other SSAs is used for predicting long-term movement and space use of animals (Potts & Schlägel 2020, Signer et al. 2017). Consequently, we encourage users aiming to make such long-term predictions to use the GF-iSSA, since the predictions that account for unobserved spatial variation are likely to have better predictive power than analyses that cannot account for this variation.

## AUTHORS’ CONTRIBUTIONS

R.A.G., F.L., S.M. and U.E.S. contributed conceptually and formulated the model. R.A.G., F.L. and S.M. provided the practical implementation of the model. T.W.G and G.A.B contributed with the data and ecological concepts of the manuscript. All authors assisted in writing and editing the manuscript.

## ACKNOWLEDGEMENTS

This project was founded by the *German Research Foundation* (DFG) Grant SCHL 2259/1-1. In addition, we thank the *Wildlife Conservation Society* for providing us wolverine data.

## AUTHOR APPROVALS

All authors have seen and approved the manuscript. In addition, this manuscript has not yet been accepted or published elsewhere.

