## Supplemental material for "Accounting for unobserved spatial variation in step selection analyses of animal movement via spatial random effects"

January 17, 2023

### SUPPLEMENTARY MATERIAL

#### A: Movement kernel

We here show how to express the movement kernel in form of an exponential function (Eqn 4 in the manuscript). Similar to Avgar et al. (2016), the movement kernel can be expressed as an exponential function as follows:

$$
\begin{aligned}
&\phi(s_{t-2}, s_{t-1}, s_t; \boldsymbol{\theta}) \frac{1}{\|s_t - s_{t-1}\|} \\
&= \frac{b^a}{\Gamma(a)} \|s_t - s_{t-1}\|^{a-1} \exp\left(-b\|s_t - s_{t-1}\|\right) \frac{\exp\left(k\cos(\alpha_{s_t,s_{t-1},s_{t-2}})\right)}{2\pi I_0(k)} \frac{1}{\|s_t - s_{t-1}\|} \\
&\propto \|s_t - s_{t-1}\|^{a-1} \exp\left(-b\|s_t - s_{t-1}\|\right) \exp\left(k\cos(\alpha_{s_t,s_{t-1},s_{t-2}})\right) \frac{1}{\|s_t - s_{t-1}\|} \\
&\propto \exp\left((a-1)\log(\|s_t - s_{t-1}\|)\right) \exp\left(-b\|s_t - s_{t-1}\|\right) \exp\left(k\cos(\alpha_{s_t,s_{t-1},s_{t-2}})\right) \exp\left(-\log(\|s_t - s_{t-1}\|)\right) \\
&= \exp\left((a-1)\log(\|s_t - s_{t-1}\|) - b\|s_t - s_{t-1}\| + k\cos(\alpha_{s_t,s_{t-1},s_{t-2}}) - \log(\|s_t - s_{t-1}\|)\right) \\
&= \exp\left((a-2)\log(\|s_t - s_{t-1}\|) - b\|s_t - s_{t-1}\| + k\cos(\alpha_{s_t,s_{t-1},s_{t-2}})\right) \\
&:= \exp\left(\beta_{p+1} log(\|s_t - s_{t-1}\|) + \beta_{p+2}\|s_t - s_{t-1}\|) + \beta_{p+3}\cos(\alpha_{s_t,s_{t-1},s_{t-2}})\right) \\
&:= \exp(\zeta(s_t, s_{t-1}, s_{t-2})),
\end{aligned}
\tag{S1}
$$

where $\Gamma(a)$ represents the Gamma and function $I_0(k)$ represents the modified Bessel function of order 0. In addition, $a$ and $b$ represent the shape and rate parameters of the Gamma distribution. The concentration parameter of the von Mises distribution is represented by $k$. Thus, after fitting the model the users can extract the movement kernel parameters directly by transforming the corresponding estimates based on Eqn S1. Consequently, we have:

- $a = \beta_{p+1} + 2$
- $b = -\beta_{p+2}$
- $k = \beta_{p+3}$

#### B: Bayesian version of the Poisson trick

As mentioned in the manuscript, Aarts et al. (2012) showed that the estimated slopes of a conditional Non-homogeneous Poisson process (NHPP) are the same of the ones using an unconditional NHPP. Here we show that this is also true for the Bayesian framework.

In Section 2.2 the observation model for the movement process was reformulated as a collection of NHPP models, with log-likelihood stated in Eqn 6 in the manuscript. In a similar context, Aarts et al. (2012), showed that the maximum likelihood estimators of the model coefficients using an unconditional NHPP with added random intercepts are equivalent to the ones from the conditional NHPP. Since we use a Bayesian framework, we need to show that this equivalence also holds for the full posterior distributions. We let the random intercepts have prior uniform densities on the interval $(-c, c)$ for some positive real number. The joint observation density for multiple unconditional NHPPs with one observed location $s_t$ at each time point $t$ is defined as

$$p(\{s_3, \ldots, s_T\}|s_1, s_2, \boldsymbol{\theta}, \{\alpha_3, \ldots, \alpha_T\}) = \prod_{t=3}^{T} \frac{\Lambda(s_t|s_{t-1}, s_{t-2}, \mathbf{X}(s_t)) \exp(\alpha_t)}{\exp\left(\int_S \Lambda(q_t|s_{t-1}, s_{t-2}, \mathbf{X}(q_t)) \exp(\alpha_t)\, \mathrm{d}q_t\right)}. \tag{S2}$$

We now let $\boldsymbol{\theta}$ denote the collection of all latent parameters in the model, and write $p(\boldsymbol{\theta})$ for their joint prior density. Multiplying Eqn (S2) with the prior densities for $\boldsymbol{\theta}$ and $\alpha_t$, and integrating over all $\alpha_t$, leads to the following expression, that is proportional to the posterior density for $\boldsymbol{\theta}$:

$$\begin{aligned}
p(\boldsymbol{\theta} \mid \text{obs}) &= \frac{p(\boldsymbol{\theta})}{p(\text{obs})} \prod_{t=3}^{T} \int_{-c}^{c} p(\alpha_t) p(s_t \mid s_{t-1}, s_{t-2}, \boldsymbol{\theta}, \alpha_t)\, \mathrm{d}\alpha_t \\
&= \frac{p(\boldsymbol{\theta})}{p(\text{obs})} \prod_{t=3}^{T} \frac{1}{2c} \int_{-c}^{c} \frac{\Lambda(s_t|s_{t-1}, s_{t-2}, \mathbf{X}(s_t)) \exp(\alpha_t)}{\exp\left(\int_S \Lambda(q_t|s_{t-1}, s_{t-2}, \mathbf{X}(q_t)) \exp(\alpha_t)\, \mathrm{d}q_t\right)}\, \mathrm{d}\alpha_t \\
&= \frac{p(\boldsymbol{\theta})}{p(\text{obs})} (2c)^{T-2} \prod_{t=3}^{T} \left[ -\frac{1}{\exp\left(\int_S \Lambda(q_t|s_{t-1}, s_{t-2}, \mathbf{X}(q_t)) \exp(\alpha_t)\, \mathrm{d}q_t\right)} \frac{\Lambda(s_t|s_{t-1}, s_{t-2}, \mathbf{X}(s_t))}{\int_S \Lambda(q_t|s_{t-1}, s_{t-2}, \mathbf{X}(q_t))\, \mathrm{d}q_t} \right]_{-c}^{c}.
\end{aligned} \tag{S3}$$

When $c \to \infty$, the limiting behaviour of Eqn (S3) is

$$(2c)^{2-T} p(\boldsymbol{\theta} \mid \text{obs}) \to \frac{p(\boldsymbol{\theta})}{p(\text{obs})} \prod_{t=3}^{T} \frac{\Lambda(s_t|s_{t-1}, s_{t-2}, \mathbf{X}(s_t))}{\int_S \Lambda(q_t|s_{t-1}, s_{t-2}, \mathbf{X}(q_t))\, \mathrm{d}q_t}. \tag{S4}$$

Thus, when $c$ is large, adding random intercepts to the model and treating it as an unconditional NHPP yields the same posterior distribution as the intended conditional NHPP, with respect to the original parameters $\boldsymbol{\theta}$. The effective joint likelihood is the joint likelihood for many conditional NHPPs, based on our Initial density from Forester et al. (2009) (Eqn 1 in the manuscript). The wide uniform distribution for the $\alpha_t$ can be replaced by a Gaussian distribution with zero mean and a large fixed variance as presented by Muff et al. (2020) to be compatible to the class of models that can be fitted with INLA.

#### C: Supplementary figures

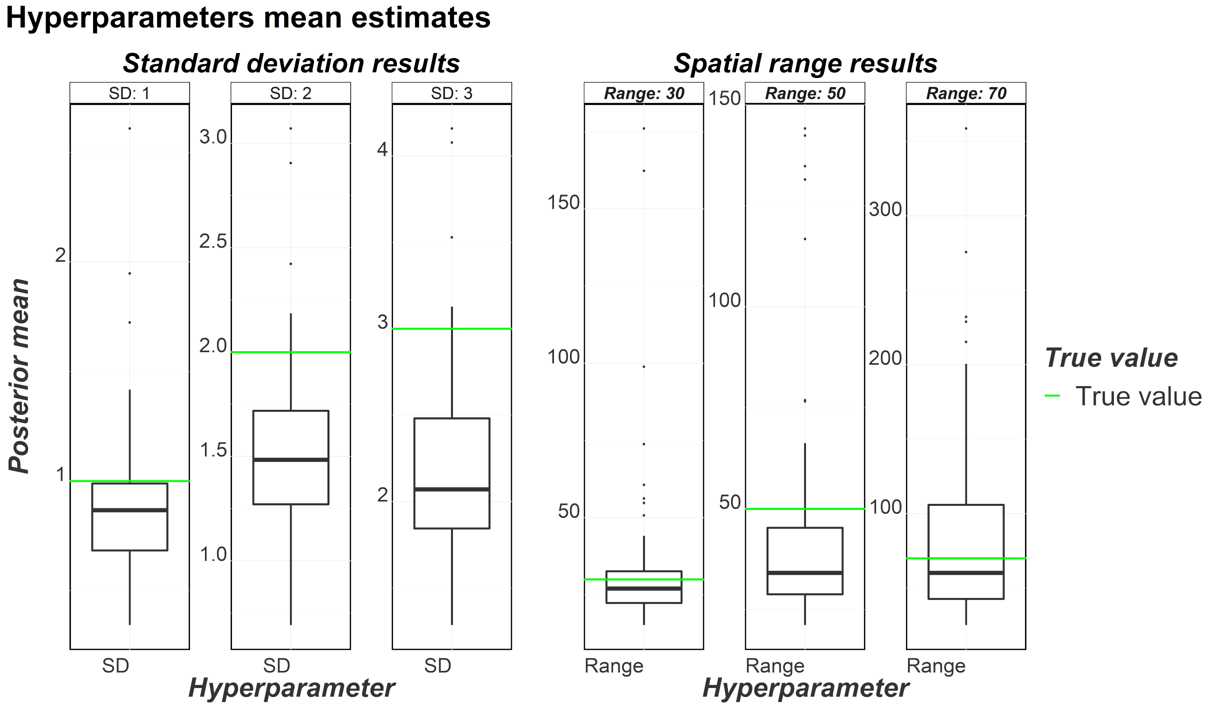


Figure S1: Gaussian field hyperparameters: Standard deviation and spatial range posterior mean estimates. Each box-plot represents the corresponding three hyperparameter combinations (75 tracks).
